# Restricted and non-essential redundancy of RNAi and piRNA pathways in mouse oocytes

**DOI:** 10.1101/678177

**Authors:** Eliska Taborska, Josef Pasulka, Radek Malik, Filip Horvat, Irena Jenickova, Zoe Jelić Matošević, Petr Svoboda

## Abstract

Germline genome defense evolves to recognize and suppress retrotransposons. One of defensive mechanisms is the PIWI-associated RNA (piRNA) pathway, which employs small RNAs for sequence-specific post-transcriptional and transcriptional repression. The loss of the piRNA pathway in mice causes male sterility while females remain fertile. Unlike spermatogenic cells, mouse oocytes have also RNA interference (RNAi), another small RNA pathway capable of retrotransposon suppression. To examine whether RNAi compensates the loss of the piRNA pathway in mouse oocytes, we produced a new RNAi pathway mutant *Dicer*^SOM^ and crossed it with a catalytically-dead mutant of *Mili*, an essential piRNA gene. Normal follicular and oocyte in double mutants showed that RNAi does not suppress a strong piRNA knock-out phenotype. However, we observed redundant and non-redundant targeting of specific retrotransposons. Intracisternal A Particle retrotransposon was mainly targeted by the piRNA pathway, MT and RLTR10 retrotransposons were targeted mainly by RNAi. Importantly, only double mutants showed increased background levels of transcripts potentially originating from intact LINE-1 elements. Our results thus show that while both small RNA pathways are simultaneously expendable defense pathways for ovarian oocyte development, yet another transcriptional silencing mechanism must mediate LINE-1 repression in female germ cells.

**Author summary:** Retrotransposons are mobile genomic parasites causing mutations. Germ cells need protection against retrotransposons to prevent heritable transmission of their new insertions. The piRNA pathway is an ancient germline defense system analogous to acquired immunity: once a retrotransposons jumps into a specific “genomic checkpoint”, it is recognized and suppressed. Remarkably, the murine piRNA pathway is essential for spermatogenesis but not oocyte development. In contrast, zebrafish lacking the piRNA pathway do not develop any germ cells. It was hypothesized that RNA interference pathway could rescue oocyte development in mice lacking the piRNA pathway. RNA interference also targets retrotransposons and is particularly enhanced in mouse oocytes. To test this hypothesis, we engineered mice lacking both pathways and observed that oocytes in these mice develop normally, which argues against the hypothesis. Furthermore, analysis of individual retrotransposon groups revealed that in specific cases the two pathways can mutually compensate each other. However, this ability is restricted to specific retrotransposon groups and seems to evolve stochastically. Finally, our results also indicate that there must be yet another layer of retrotransposon silencing in mouse oocytes, which prevents high retrotransposon activity in the absence of piRNA and RNA interference pathways.

## INTRODUCTION

Genome integrity in the germline is important for intact transmission of genetic information into progeny. However, it is being disturbed by retrotransposons, mobile genomic parasites reproducing through a “copy & paste” strategy (reviewed in [1]). Although retrotransposons threaten genome integrity, they occasionally also provide new functional gene elements, such as promoters, enhancers, exons, splice junctions, or polyA signals, thus contributing to evolution of new traits (reviewed in [1]). Retrotransposons can be categorized according to the presence of long terminal repeats (LTRs) and retrotransposition autonomy [2]. While contributions of different families vary, their collective contribution to the mammalian genome content is substantial: annotatable retrotransposon insertions comprise about a half of human and mouse genomes [3, 4]. The mouse genome hosts two actively retrotransposing autonomous retrotransposons: non-LTR element LINE-1 (L1) and LTR element Intracisternal A Particle (IAP) [5–7]. L1 (reviewed in [8]) is the most successful mammalian autonomous retrotransposon with 868,000 insertions in the mouse genome [4]. L1 has several unique adaptations, such as a unique complex bidirectional promoter [9, 10] and retrotransposition in *cis* [11]. IAP is an endogenous retrovirus, which invaded the mouse genome relatively recently [6]. RepeatMasker identifies approximately 13 000 IAP insertions, of which 44% represent solo LTRs [12].

Various transcriptional and post-transcriptional mechanisms evolved to repress retrotransposons (reviewed in [13]). A key mechanism suppressing retrotransposons in the mammalian germline is the PIWI-interacting RNA (piRNA) pathway, which combines post-transcriptional and transcriptional silencing (reviewed in [14]). The piRNA pathway relies on specific genomic “checkpoints” for invading mobile elements regions (piRNA clusters) that give rise to piRNAs, 25-30 nucleotides long RNAs loaded onto PIWI subgroup of the Argonaute protein family, which guide retrotransposon recognition and repression [15]. Complex piRNA system starts with processing of primary transcripts from piRNA clusters into piRNAs, which guide cleavage of retrotransposon transcripts. This triggers production of a secondary piRNA pool, which further facilitate processing of primary transcripts into piRNAs (reviewed in [14]).

Mice use three PIWI proteins PIWIL1 (MIWI), PIWIL2 (MILI), and PIWIL4 (MIWI2). PIWIL3, the fourth mammalian PIWI protein, was found in bovine oocytes [16] but *Piwil3* gene was apparently lost in the common ancestor of mice and rats. All three mouse PIWI proteins are essential for spermatogenesis but not oogenesis [17–19] although the piRNA pathway operates during oogenesis where it targets retrotransposon expression [20, 21]. MILI is a cytoplasmic protein, which generates primary piRNAs and secondary piRNAs by so-called ping-pong mechanism with itself or with MIWI2 [22–24]. MIWI2 shuttles to the nucleus and mediates transcriptional silencing through DNA and histone methylation of retrotransposon loci [25, 26]. In males, MIWI2 expression ceases around birth whereas MILI is important for clearance of retrotransposon transcripts in postnatal testes [24, 27]. MILI and MIWI2 cooperate on silencing of L1 and IAP (non-LTR and LTR) retrotransposons in mouse fetal testes [28, 29].

It is unknown what accounts for the strikingly different phenotypes of piRNA pathway mutants in murine male and female germlines. In contrast to mice, *Drosophila* or zebrafish females lacking piRNA pathway components are sterile [17–19, 30]. It was hypothesized that the loss of the piRNA pathway in the mouse female germline could be compensated by RNA interference (RNAi) [31, 32]. Canonical RNAi is defined as sequence-specific RNA degradation induced by long double-stranded RNA (dsRNA) [33] and is active in mouse oocytes [34, 35]. The canonical RNAi starts with processing of long dsRNA by RNase III Dicer into ∼22 nucleotides long small interfering RNAs (siRNAs), which guide sequence-specific cleavage of perfectly complementary RNAs by Argonaute 2 (AGO2) (reviewed in [36]). Endogenous RNAi in mouse oocytes and zygotes was shown to target mobile elements and regulate gene expression [31, 32, 37–40]. For example, L1 can be targeted in oocytes by piRNAs and endo-siRNAs [32, 40–43]. Analysis of small RNA-seq data from mouse oocytes [43] shows that piRNA and RNAi pathways may target retrotransposons simultaneously but the extent of repression for individual retrotransposons by each pathway could vary (Fig. 1). Importantly, endogenous RNAi in mouse oocytes evolved through exaptation of an MT retrotransposon insertion in intron 6, which functions as an oocyte-specific promoter producing a unique truncated Dicer isoform (denoted Dicer^O^). *Dicer*^O^ transcripts accumulate during oocyte’s growth and persist into the zygote [44]. Based on RNA sequencing (RNA-seq), *Dicer*^O^ is expressed already in non-growing oocytes 5 days postpartum (5dpp) [45].

**Figure 1.**
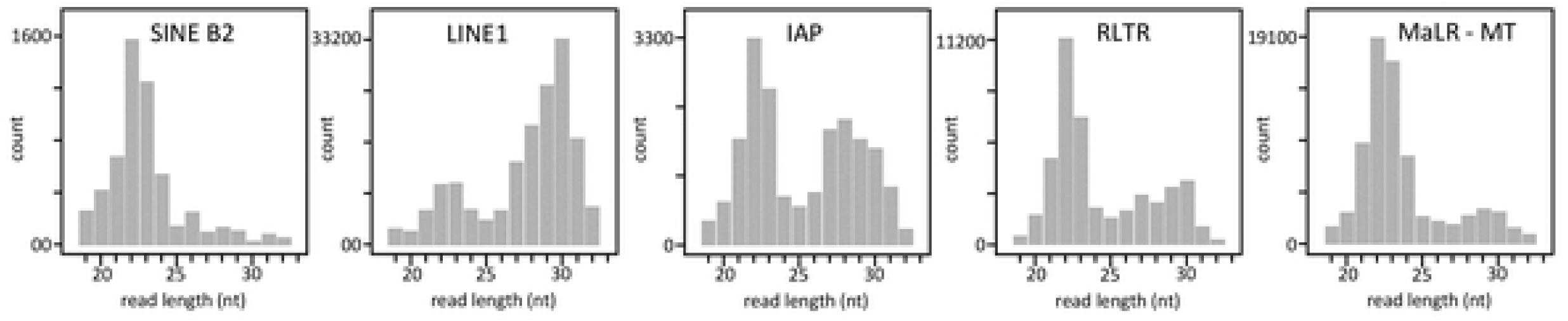
Different retrotransposons are associated with different length distribution of small RNAs from mouse oocytes suggesting partially redundant repression of retrotransposons by endogenous RNAi and piRNA pathways. Based on RNA-seq data fromYang et al., 2016 [43].

To address the significance of redundancy of RNAi and piRNA pathways, we produced and analyzed mice lacking both pathways. We first produced a novel mouse model lacking *Dicer*^O^ (denoted *Dicer*^SOM^) and then crossed it with *Mili*^DAH^ mice expressing catalytically inactive MILI [24]. We show that engagement of piRNA and RNAi pathway in repression of retrotransposons is stochastic and occasionally synergic. The simultaneous loss of RNAi and piRNA pathways uncovers redundant targeting of L1 and extended L1 derepression but does not affect oocyte development. Thus, while piRNAs are essential for spermatogenesis and show restricted redundancy with RNAi, they do not have essential role in ovarian oocyte development that would be masked by RNAi.

## RESULTS

### Generation of Dicer^SOM^ mice – a novel RNAi-deficient model

The oocyte-specific Dicer^O^ isoform comprises the majority of Dicer protein in mouse oocytes and deletion of its LTR-derived promoter (*Dicer*^ΔMT^ mutant) phenocopies conditional *Dicer* knock-out in mouse oocytes [38, 44]. However, we have discovered that there is a second LTR insertion (MTA) in intron 6 (Fig. 2A), which can also produce a truncated Dicer variant and reduce phenotype penetrance in *Dicer*^ΔMT/ΔMT^ mice [46]. Thus, to eliminate truncated Dicer expression in mouse oocytes, we generated another modified *Dicer* allele (denoted *Dicer*^SOM^), which has an HA-tag at the N-terminus and lacks introns 2-6 (Fig. 2A). It was shown that N-terminal HA-tag on Dicer does not produce a phenotype and allows for its detection with highly-specific anti-HA antibodies [47].

**Figure 2.**
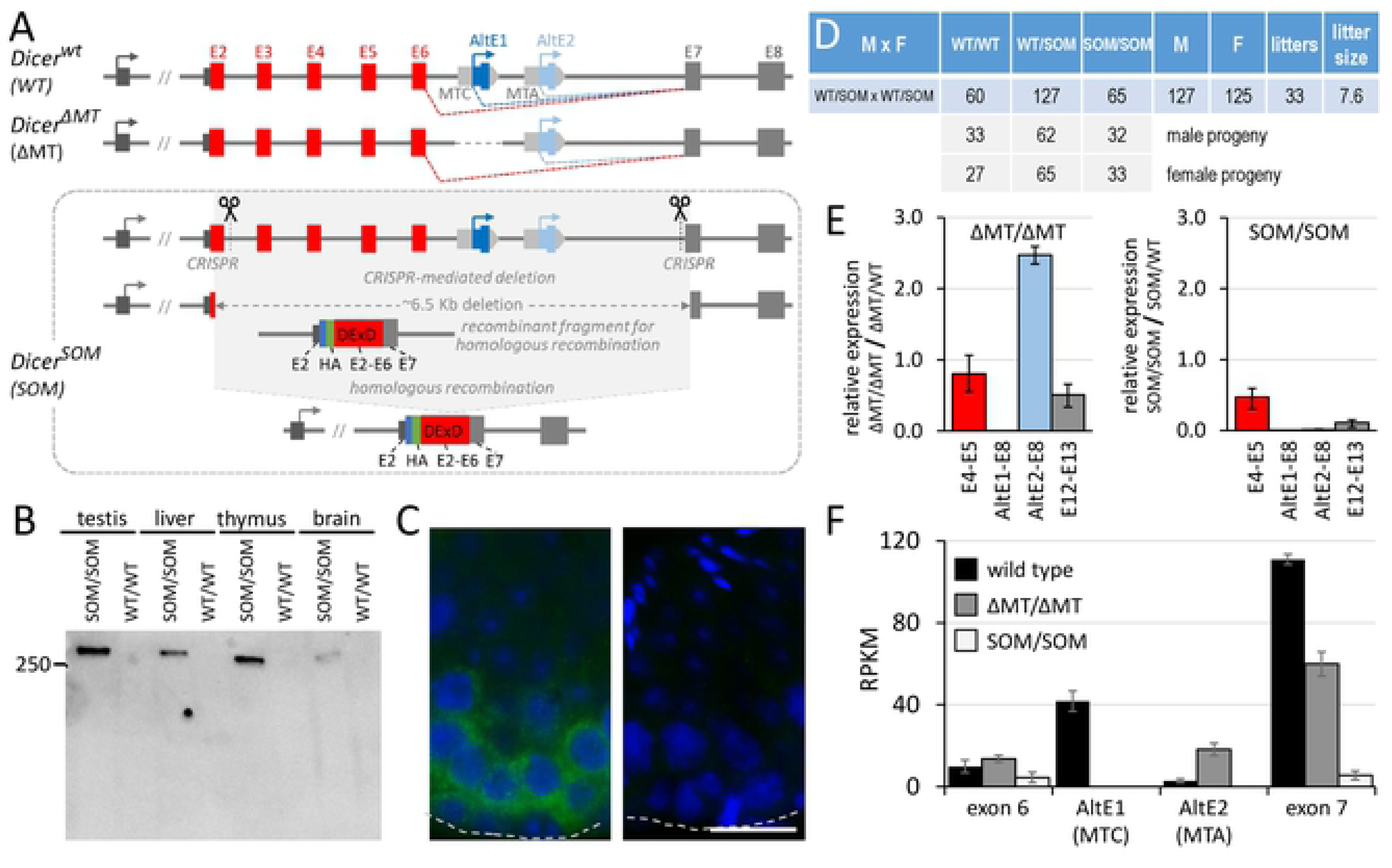
*Dicer*^SOM^ mouse model. (A) Schematic depiction of the 5’ gene structure of *Dicer* and *Dicer*^y^and *Dicer*^SOM^ models. *Dicer*^SOM^ model was produced by removing ∼6.5 kb of genomic DNA using CRISPR nucleases followed by homologous recombination with a construct encoding an HA-tag and exons 2-7. The resulting allele lacks introns 2 to 6 while encoding HA-tagged full-length Dicer. (B) *Dicer*^SOM^ mice express HA-tagged full-length Dicer. 40 μg of total protein lysate were loaded per lane. (C) Dicer^SOM^ protein has cytoplasmic expression. Shown are sections of seminiferous tubules with high Dicer^SOM^ signal in spermatogenic cells of *Dicer*^SOM/SOM^ mice. Dashed lines indicate positions of basal lamina. (D) *Dicer*^SOM/SOM^ animals are born in a Mendelian ratio upon crossing *Dicer*^SOM/wt^ parents. (E) Analysis of Dicer expression by qPCR in oocytes of homozygous *Dicer*^ΔMT^ and *Dicer*^SOM^ mutants. Colors indicate cDNA region amplified by primers localized in exons as described in the x-axis legend. *Dicer*^ΔMT/ΔMT^ mice show loss of expression of the MTD-driven Dicer variant and a relative increase of expression from the downstream MTA LTR insertion. *Dicer*^SOM/SOM^ mice show complete loss of short Dicer isoform. Data were normalized to oocytes from heterozygous littermates. (F) Analysis of Dicer expression in oocytes of wild type animals and homozygous *Dicer*^ΔMT^ and *Dicer*^SOM^ mutants by RNA-seq. Depicted are RPKM values per exon.

*Dicer*^SOM^ allele was generated in mouse embryonic stem cells (ESCs) by cleaving intron 2 and intron 6 with CRISPR nucleases followed by homologous recombination with a construct carrying HA-tag fused to the 5’ end of *Dicer* cDNA containing coding sequence of exon 2 to exon 7. Homologous recombination arms contained intron 1 and intron 7 (Fig. 2A). An ESC line carrying the desired recombination event (Fig. S1) was subsequently used to make mouse chimaeras from which we established a mouse line. Analysis of *Dicer*^SOM^ mice showed that the full-length HA-tagged Dicer could be detected by Western blotting in organ lysates (Fig. 2B) and by immunofluorescent staining of histological sections albeit the signal was relatively low with the protocol and antibody used (Fig. 2C). Breeding of *Dicer*^SOM/wt^ animals yielded progeny with *Dicer*^SOM/SOM^ genotypes at the expected Mendelian ratio (Fig. 2D). *Dicer*^SOM/SOM^ mice appeared normal and males were fertile suggesting that the introduced modification did not perturb normal full-length Dicer function. Importantly, *Dicer*^SOM/SOM^ females were sterile, which was the expected phenotype caused by the lack of Dicer^O^. Analysis of *Dicer*^SOM/SOM^ oocytes by qPCR and RNA-sequencing (RNA-seq) showed the loss of *Dicer*^O^-encoding transcript and minimal levels of the full-length Dicer isoform mRNA, which contrasted with *Dicer*^ΔMT/ΔMT^ oocytes where expression from the MTA LTR was enhanced (Fig. 2E, F). Thus, *Dicer*^SOM/SOM^ mice solved the problem with residual Dicer^O^ expression observed in *Dicer*^ΔMT/ΔMT^ mice.

### *Dicer*^SOM^ mutants phenocopy oocyte-specific *Dicer* knock-out

Next, we analyzed the phenotype of *Dicer*^SOM/SOM^ oocytes and compared that with known phenotypes of *Dicer* and *Ago2* mutants [31, 38, 44]. Fully-grown *Dicer*^SOM/SOM^ oocytes showed high frequency of meiotic spindle defects upon resumption of meiosis (Fig. 3A, B) and upregulated mRNA levels of RNAi targets (Fig. 3C) consistent with RNAi pathway deficiency. For precise characterization of transcriptome changes in *Dicer*^SOM/SOM^ oocytes, we performed RNA-seq of fully-grown germinal vesicle-intact (GV) oocytes. When comparing mRNA changes in *Dicer*^ΔMT/ΔMT^ and *Dicer*^SOM/SOM^ oocytes, the later showed more deregulated transcriptome (Fig. 3D). In addition, we observed upregulated mRNAs of many genes, which would have potential to be targeted by RNAi (Fig. 3D, red points), i.e. genes where complementary endo-siRNAs were identified in RNA-seq datasets from mouse oocytes [32, 41, 43]. This was consistent with the principal component analysis (PCA) where *Dicer*^ΔMT/ΔMT^ samples localized between wild type and *Dicer*^SOM/SOM^ samples (Fig. 3E). We included into PCA also transcriptomes of previously analyzed *Dicer* and *Ago2* mutants [31]. Although the experimental variability (“bench effect”) separated our and other datasets along the PC1 axis, the data showed an apparent separation of wild type and RNAi mutant samples along the PC2 axis. *Dicer*^SOM/SOM^ transcriptome appeared to be more distant from wild type samples than *Dicer*^ΔMT/ΔMT^ transcriptome and in a parallel position with Stein et al. mutants [31] relative to wild type samples (Fig. 3E).

**Figure 3.**
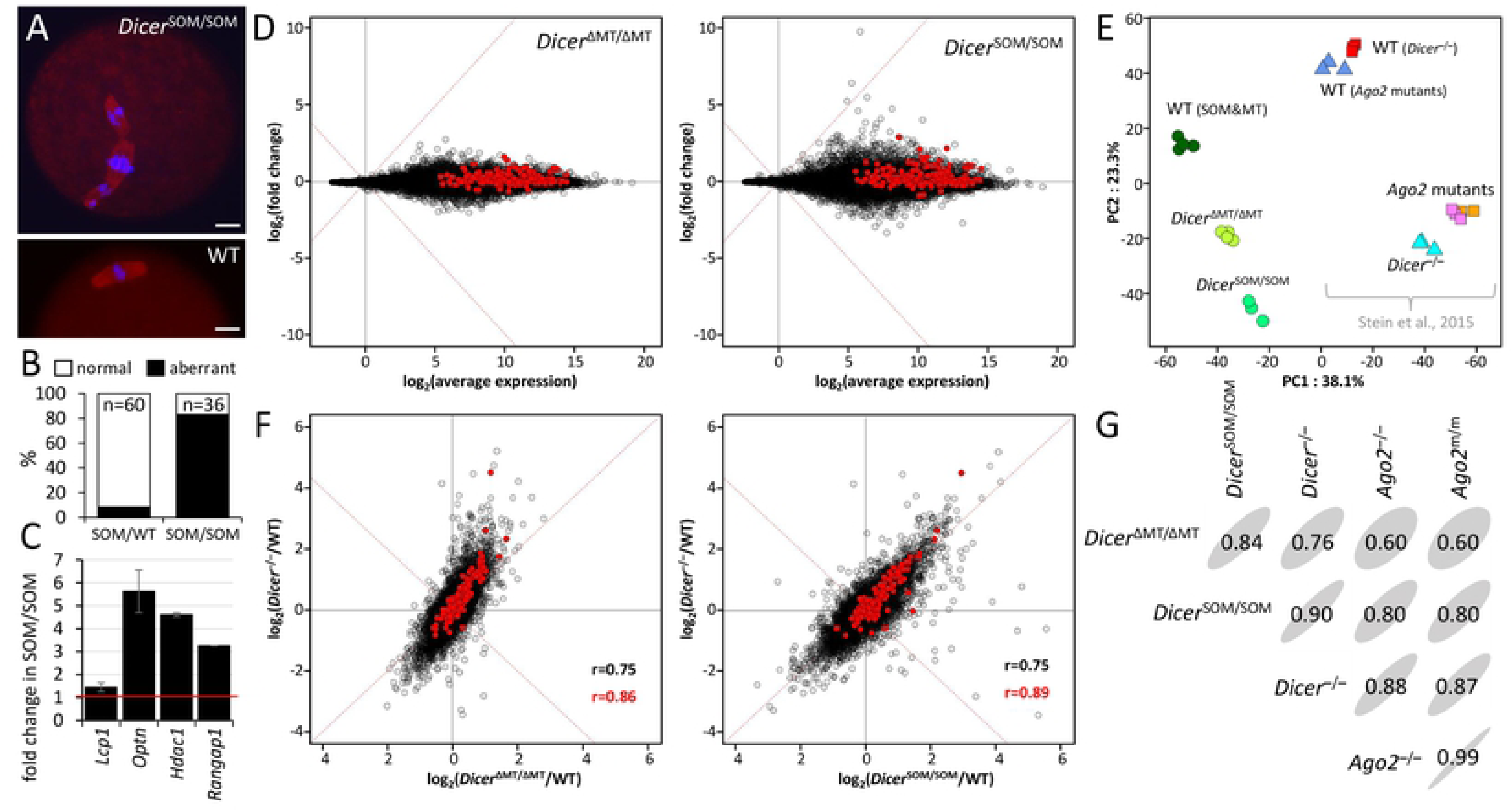
*Dicer*^SOM/SOM^ animals phenocopy oocyte-specific Dicer knock-out. (A) Oocytes of *Dicer*^SOM/SOM^ mice exhibit spindle defects like *Dicer*^−/−^ or *Dicer*^ΔMT/ΔMT^ oocyte [38, 39, 44]. Size bar = 10 µm. (B) Frequency of meiotic spindle defects. (C) mRNA expression of selected RNAi targets analyzed by qPCR. (D) MA plots depicting transcriptome changes in *Dicer*^ΔMT/ΔMT^ and *Dicer*^SOM/SOM^ oocytes relative to wild type oocytes. (E) PCA analysis of RPKM transcriptome changes in oocytes of different mutants, which includes previously published data from Dicer and Ago mutants. (F) Comparison of mRNA expression changes in *Dicer*^ΔMT/ΔMT^ and *Dicer^S^*^OM/SOM^ oocytes (x-axis) with mRNA changes in oocytes with conditional Dicer knock-out (y-axis). (G) Correlation matrices of retrotransposon-derived transcriptome in different mutant oocytes. Elliptic shapes reflect sizes of inscribed correlation coefficients for easier visual navigation.

When comparing relative transcriptome changes of Dicer mutants to accompanying wild type controls (Fig. 3F, S2), *Dicer*^ΔMT/ΔMT^ and *Dicer*^SOM/SOM^ showed good overall correlation with transcriptome changes in *Dicer*^−/−^ oocytes (r=0.75) and relative changes of predicted RNAi targets correlated slightly better in *Dicer*^SOM/SOM^ (r=0.89) than changes in *Dicer*^ΔMT/ΔMT^ (r=0.86) oocytes. While the difference in correlation coefficient is minimal, putative RNAi targets and other upregulated genes tend to be distributed above the diagonal when comparing *Dicer^ΔMT/ΔMT^* and *Dicer^−/−^* samples and more along it when comparing *Dicer*^SOM/SOM^ with *Dicer*^−/−^ (Fig. 3F). This means that transcriptome changes in *Dicer*^SOM/SOM^ better mimic changes in *Dicer^−/−^* than transcriptome changes in *Dicer*^ΔMT/ΔMT^ and show that *Dicer*^SOM/SOM^ model is superior to *Dicer*^ΔMT/ΔMT^ as oocyte-specific RNAi deficiency model. As RNAi also targets repetitive sequences, we analyzed correlations of relative changes of retrotransposon RNAs in our datasets and *Dicer* and *Ago2* mutants [31]. In this case, *Dicer*^SOM/SOM^ mutants correlated with *Dicer^−/−^* mutant (r=0.90) even better than *Dicer^−/−^* mutant with *Ago2* mutants (Fig. 3G), which is remarkable considering entirely independent experimental analysis. Since *Dicer*^SOM/SOM^ females closely phenocopy conditional *Dicer* knock-out in oocytes, the *Dicer*^SOM^ model is excellent for studying simultaneous loss of RNAi and piRNA pathways in oocytes because it overcomes arduous breeding of the *Mili*^DAH^ model with a conditional *Dicer* knock-out.

### RNAi & piRNA mutant females show no signs of disturbed ovarian development of oocytes

Mouse mutants lacking piRNA and RNAi pathways in mouse oocytes should reveal whether or not mutants of single pathways manifest the full extent of small RNA-mediated retrotransposon repression because both pathways can suppress mobile elements. Accordingly, we crossed *Dicer*^SOM^ mice with *Mili*^DAH^ mice carrying a mutation in the DDH catalytic triad of MILI where the second aspartic acid is mutated to an alanine [24]. As *Mili*^DAH/DAH^ males and *Dicer*^SOM/SOM^ females are sterile, we first crossed *Mili*^DAH/DAH^ females to *Dicer*^SOM/SOM^ males (both on C57Bl/6 background) and then crossed their double heterozygote progeny to obtain *Dicer*^SOM/WT^, *Mili*^DAH/DAH^ females and Dicer^SOM/SOM^, *Mili*^DAH/WT^ males, which were crossed to obtain females lacking in oocytes both pathways (*Dicer^SOM/SOM^*, *Mili*^DAH/DAH^, “double KO”), only RNAi (*Dicer*^SOM/SOM^, *Mili*^DAH/WT^), only piRNA (*Dicer*^SOM/WT^, *Mili*^DAH/DAH^), or none (*Dicer*^SOM/WT^, *Mili*^DAH/WT^). Importantly, phenotype analysis of double mutants was restricted to ovarian phenotypes manifested until the fully-grown GV oocyte stage because the loss of RNAi causes meiotic spindle defects in ovulated oocytes and impairs further development [38, 39, 44].

Females of the four above-described genotypes of age between 8 and 18 weeks were analyzed for ovarian weight and histology and number of fully-grown GV oocytes recovered from non-superovulated animals (Fig. 4A-C). Double KO females showed normal ovarian morphology and weight of ovaries (Fig. 4A, B). The presence of antral follicles and *corpora lutea* in ovaries of double KO females suggested that the loss of RNAi on the piRNA knock-out background does not reveal a strong ovarian phenotype. This was further supported by recovering normal number of fully-grown oocytes from double KO ovaries (Fig. 4C). We also did not observe increased DNA damage in oocytes (data not shown). Elimination of both pathways thus did not affect follicular growth and ovarian oocyte development, which shows that RNAi pathway does not mask an essential function of piRNAs during ovarian oogenesis.

**Figure 4.**
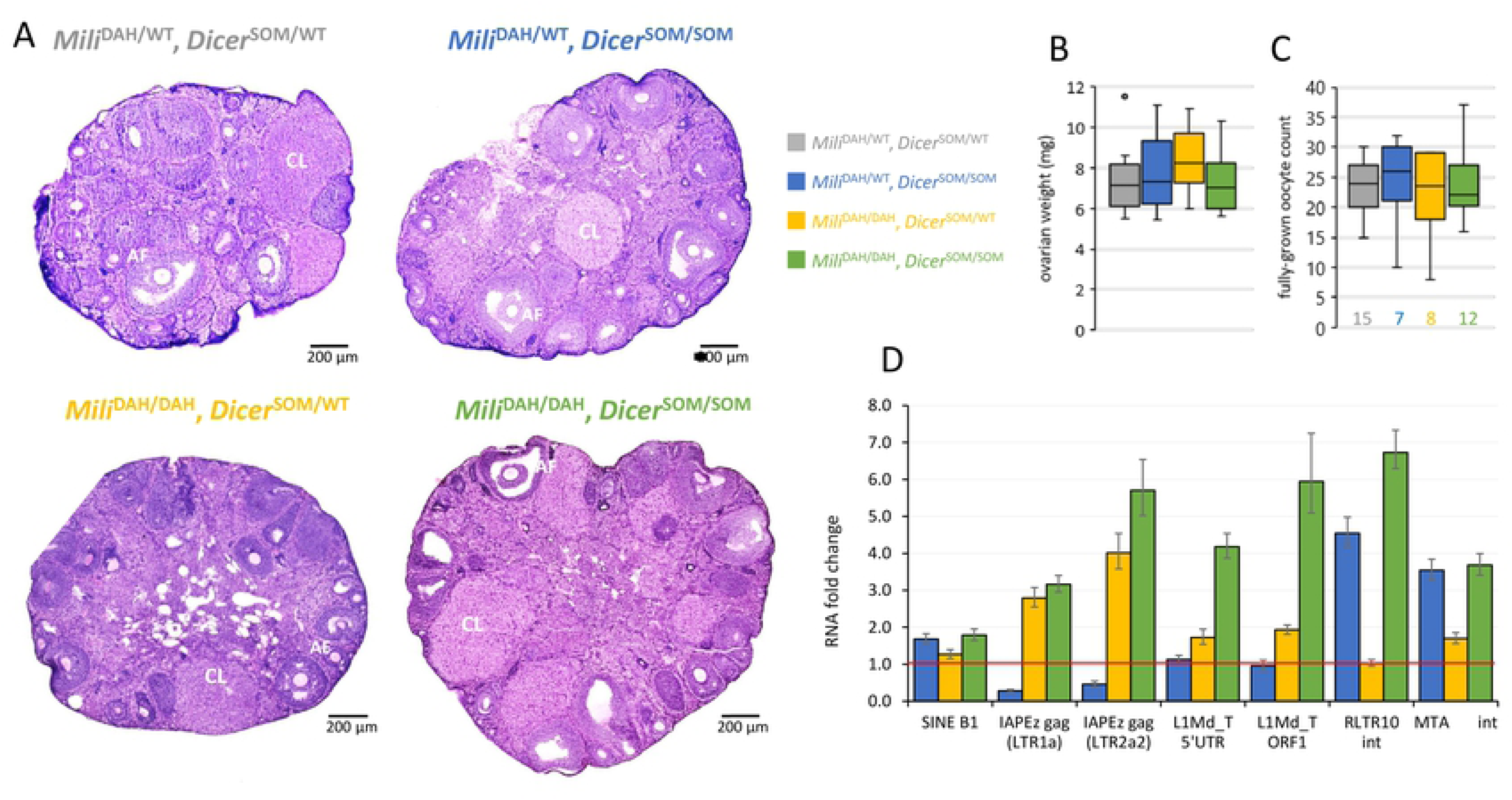
RNAi (*Dicer*^SOM/SOM^) and piRNA (*Mili*^DAH/DAH^) knock-out analysis. (A) Representative histological sections of ovaries stained with hematoxylin and eosin. Abbreviations: CL – *corpus luteum*, AF – antral follicle. (B) Ovarian weight. Ten ovaries per genotype were analyzed for this boxplot. (C) Yield of fully-grown germinal vesicle (GV)-intact oocytes isolated from ovaries per female without superovulation. Colored numbers below boxplots indicate numbers of females with each genotype. (E) qPCR analysis of retrotransposon expression in mutant oocytes. Error bar = SD.

### Varying redundancy of retrotransposon repression by RNAi and piRNA pathways

To complete the analysis of double mutants, we examined retrotransposon transcript levels. Previous studies showed upregulation of specific retrotransposon types in single pathway mutants but the extent to which retrotransposons are simultaneously regulated and how one pathway could compensate for the loss of the other one was not clear. We analyzed by qPCR active subfamilies of autonomous retrotransposon L1 (L1Md_T) and IAP (IAPEz) and selected non-autonomous elements SINE B1, RLTR10, and MTA. Comparison of oocytes of the four genotypes (double KO, RNAi KO, piRNA KO, and double heterozygote) showed interesting diversity of transcript level changes (Fig. 4D).

IAP seemed to be targeted by the piRNA pathway and was even better suppressed in RNAi knock-out oocytes. Levels of IAPs internal sequence showed low if any increase when RNAi was lost in addition to the piRNA pathway (Fig. 4D). A strong effect of double knock-out was observed on L1Md_T expression. While RNAi knock-out had no effect and *Mili* knock-out showed only slightly increased L1 expression, the loss of both pathways caused about six-fold increase in expression suggesting that both pathways target L1 and can mutually compensate loss of a single pathway. In contrast, RNAi primarily targeted RLTR10. *Mili* knock-out had no effect on RLTR10 expression on its own but had minor additive effect on *Dicer*^SOM/SOM^ background (Fig. 4D). These data demonstrate that RNAi and piRNA pathways suppress some but not all retrotransposons in a redundant manner and can partially compensate for each other’s loss. However, even if both pathways are eliminated and fail to suppress retrotransposons in non-growing oocytes or during oocyte growth, it does not lead to substantial retrotransposon mobilization and failure of ovarian oogenesis.

## DISCUSSION

We experimentally tested whether or not RNAi acts redundantly and compensates the loss of piRNAs in the murine female germline. Existence of such a compensation was proposed previously [31, 32] and seemed plausible given normal fertility of female mutants of the piRNA pathway [17–19, 30] and simultaneous presence of RNAi and piRNA pathways in the oocyte [32, 40]. Furthermore, the vertebrate female germline is not insensitive to the loss of the piRNA pathway *per se* as shown in zebrafish [48]. To test the compensation hypothesis, we engineered a novel mouse mutant *Dicer*^SOM^, which could be directly crossed with a piRNA pathway mutant to obtain females deficient in both pathways in the germline in a simple crossing scheme.

*Dicer*^SOM^ mice express a tagged Dicer protein and lack expression of the short Dicer^O^ variants, which support RNAi in oocytes [46]. The *Dicer*^SOM^ model is superior to our earlier *Dicer*^ΔMT^ RNAi-deficient mouse model that lacked the main *Dicer*^O^ promoter and phenocopied oocyte-specific *Dicer* knock-out phenotype [44]. *Dicer*^ΔMT^ model was problematic because of varying penetrance of the phenotype due to an alternative LTR promoter that could also drive Dicer^O^ expression [46]. *Dicer*^SOM^ model faithfully phenocopies the loss of Dicer in mouse oocytes including derepression of RNAi-targeted genes and retrotransposons. Low levels of the full-length Dicer remain expressed in *Dicer*^SOM/SOM^ oocytes (Fig. 2F). The residual full-length Dicer is able to sustain miRNA biogenesis (data not shown) but not efficient RNAi, as shown by transcriptome remodeling and phenocopy of Dicer null phenotype. This is also consistent with minimal siRNA levels produced by the full length Dicer in somatic cells from dsRNA substrates, which are capable of inducing RNAi in oocytes [49, 50].

For a piRNA pathway mutant, we have chosen *Mili*^DAH^ mouse model expressing catalytically dead MILI [24]. MILI is involved in processing of primary piRNA transcripts [51]; its prenatal or postnatal loss derails the piRNA pathway and causes male but not female sterility [24, 52]. When we crossed *Dicer*^SOM^ mice with *Mili*^DAH^ mice, the double mutants revealed no effect on ovarian oogenesis, up to the diplotene stage of the first meiotic division, in which oocytes remain arrested until resumption of meiosis (reviewed in [53]). This shows that piRNA and RNAi pathways in the mouse female germline do not suppress an acute threat, which would manifest as an ovarian phenotype when both pathways are removed. However, long term consequences could be still sufficient for maintaining both pathways present through eons.

Importantly, we observed retrotransposon-specific extent of redundant repression by RNAi and piRNA pathway. We observed that both pathways suppress retrotransposons in oocytes in a partially redundant manner, which could be explained by the stochastic nature of evolution of substrates giving rise to siRNAs and piRNAs. The piRNA pathway is an adaptive defense, where the recognition of an invading retrotransposon is stochastic; it requires retrotransposon insertion into one of the piRNA clusters yielding piRNA production and retrotransposon repression [54]. The post-transcriptionally acting RNAi also evolves stochastically in a different way. RNAi requires dsRNA, which can form when retrotransposon insertions form a transcribed inverted repeat or are transcribed antisense to produce RNAs basepairing with retrotransposon transcripts. Furthermore, some retrotransposons have intrinsic potential to produce dsRNA. In particular, L1 carries an antisense promoter in its 5’UTR and can produce sense and antisense RNAs with potential to basepair and trigger RNAi (reviewed in [55]). Notably, the antisense transcription from L1 5’UTR, which exists across mammals, may represent L1’s own strategy for maintaining minimal expression levels sufficient for L1 retrotransposition in *cis* [11]; natural selection would otherwise yield L1s lacking the antisense promoter if it would significantly reduce L1’s fitness.

The lack of sterile phenotype in female mice upon removal of two pathways targeting retrotransposons could stem from specific behavior/repression of DNA-damaging retrotransposons in the mouse female germline, which would make it insensitive to the loss the piRNA pathway. Expression of retrotransposons can differ between male and female germlines as retrotransposons adapt their transcriptional control to transcription factors regulating gene expression during different parts of the germline cycle. It can be observed in RNA-seq data [45, 56–58]. Mouse oocytes have low expression of L1 and IAP (Fig. 5A). Mouse genome contains several L1 families, of which LINE1_Md_A, LINE1_Md_T(-F) and LINE1_Md_G(-F) are the most recent and contain full-length retrotransposition-competent LINE1 copies [59–61]. We examined expression of all full-length L1 elements with intact ORF in the reference C57Bl/6 genome in fully-grown oocytes by mapping our C57Bl/6 RNA-seq libraries directly on these elements (using RNA-seq of wild type oocytes from this work and from [62]). Perfectly mapping 50 nucleotide single end sequencing (50SE) showed low abundance of reads for the most expressed L1 elements from different families (Fig. 5B). A more precise analysis using perfectly-mapping reads from 125 nucleotide paired end sequencing (125PE) showed minimal if any expression of intact L1 elements with apparent signal in the 5’UTR, which seems to be associated with the antisense promoter (Fig. 5B). Since 90% of reads in 125PE libraries of fully-grown oocyte are mapable with perfect complementarity on the C57Bl/6 genome, these data suggest that most retrotransposition-competent L1 elements are transcriptionally silent in fully grown oocytes. Thus, the up to 6-fold increase in L1 RNA levels estimated by qPCR in fully-grown double mutant oocyte (Fig. 4D) is unlikely to represent significant L1 mobilization. In contrast, a tractable engineered L1 retrotransposon in *Mov10l1*^−/−^ testes showed ∼1,400-fold increase in RNA expression and 70-fold increase in retrotransposition [63].

**Figure 5.**
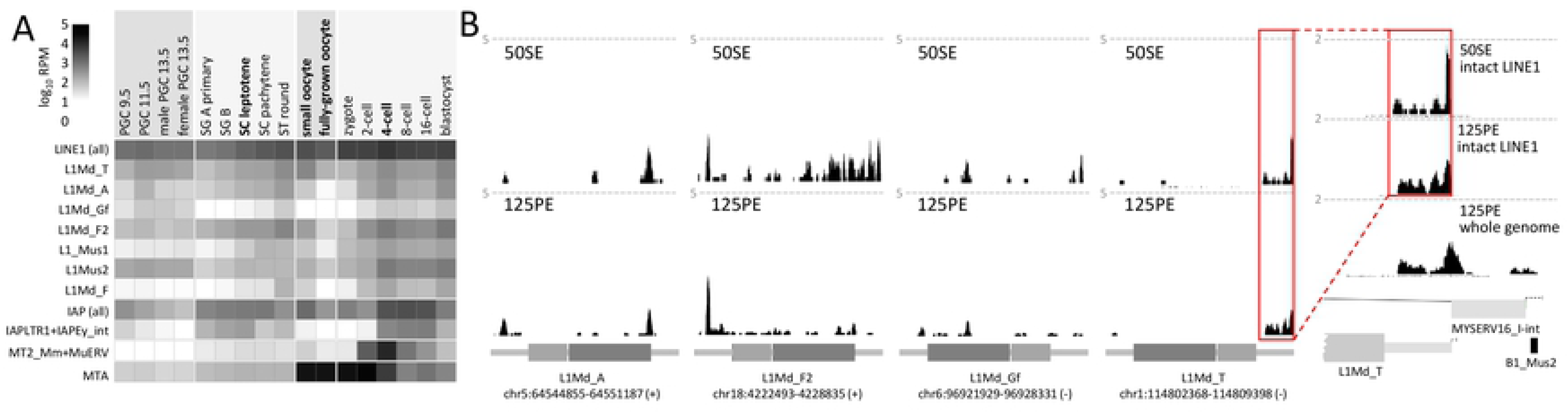
Retrotransposon expression in the female germline. (A) Relative retrotransposon expression during the germline cycle. The heatmap was produced as described previously [46] using published expression data [45, 56–58]. Depicted L1 and IAP retrotransposon subfamilies have full-length intact retrotransposon insertions in C57Bl/6 genome. (B) Most expressed intact L1 retrotransposons from the four most abundant subfamilies. Shown are UCSC genome browser snapshots of specific L1 insertions with the highest numbers of perfectly matching reads from fully-grown GV oocyte RNA-seq data: 50 nucleotide single-end (50SE) sequencing from Abe et al. [73] and 125 nucleotide paired-end (125PE) sequencing from Horvat et al. [62]. Note the reduced signal in more specific 125PE mapping in L1Md_F2. The dashed line depict 5 counts per million. L1 scheme depicts orientation of the element with ORF1 and ORF2 depicted as lighter and darker grey rectangle, respectively. L1Md_T expression likely comes most from activity of the antisense L1 promoter as the signal continues into the flanking sequence (right expanded inset).

It could be that L1 is not active during oocyte development because transcription factors controlling its transcription are present during different parts of the germline cycle. However, this would be inconsistent with the proposed role of L1 during fetal oocyte attrition, which requires variable L1 expression [64]. Thus, it is more plausible that L1 is transcriptionally repressed in the female germline by some piRNA-independent mechanism. One of the candidate transcriptional silencing mechanisms, which can evolve and act in parallel to the piRNA pathway are Krüppel-associated box domain zinc finger proteins (KRAB-ZFPs, reviewed in [65]), whose C-terminal arrays of DNA binding zinc fingers have potential to evolve into specific retrotransposon repressors. Their cofactor, TRIM28 was found enriched on some L1 subfamilies, particularly L1MdF and L1MdF2 [66]. Furthermore, KRAB-ZFP *Gm6871* was proposed to selectively target L1MdF2 elements in mice [66]. *Trim28* and *Gm6871* are expressed in 5dpp oocytes [45] making the KRAB-ZFP system an excellent candidate for a parallel functional protection of the mouse female germline, which prevents mobilization of L1 elements in the absence of RNAi and piRNA pathways. In any case, while there is no conclusive answer to the question why is the female germline insensitive to the loss of the piRNA pathway, our results rule out RNAi as a responsible mechanism and indicate that additional transcriptional silencing is involved in retrotransposon repression during oocyte development.

## MATERIALS AND METHODS

### Animals

Animal experiments were approved by the Institutional Animal Use and Care Committee (approval no. 34-2014) and were carried out in accordance with the law. *Mili*^DAH^ mice were kindly provided by Donal O’Carroll; this strain was characterized previously [24]. *Dicer*^SOM^ mice were produced in the Transgenic Unit of the Institute of Molecular Genetics ASCR, Czech Centre for Phenogenomics.

For genotyping, tail biopsies were lysed in PCR friendly lysis buffer with 0.6 U/sample Proteinase K (Thermo Scientific) at 55 °C, with 900 rpm shaking until dissolved (approx. 2.5 h). Samples were heat-inactivated at 75 °C, 15 min. The lysate was diluted three times and 1 μl used with HighQu DNA polymerase (0.5 U/reaction) master mix for PCR. Genotyping primers are provided in Table S1. PCR program was as follows – 95 °C, 3 min, 38 x [95 °C, 30 s, 58 °C, 30 s, 72 °C, 1 min], 72 °C, 3 min. PCR products were analyzed using a 1.5% agarose gel electrophoresis.

### Generation of Dicer^SOM^ mice

We first produced ESCs with the *Dicer*^SOM^ allele and then used those for producing chimeric mice and establishing *Dicer*^SOM^ line upon germline transmission of the *Dicer*^SOM^ allele. *Dicer*^SOM^ allele in ESCs [67] was generated by using CRISPR-Cas9 [68] mediated modification of the endogenous *Dicer* locus. Pairs of sgRNAs were designed to cleave *Dicer* genomic sequence in intron 2 (sequence of DNA targets: mDcr_i2a 5’-GTACCCAAATGGATAGAA-3’, mDcr_i2b 5’-GTTGGGATGGAGGTTGTT-3’) and intron 6 (sequence of DNA targets: mDcr_i6a 5’-ACTACGCTAGGTGTAAACAG-3’, mDcr_i6b 5’-TGCAGTCCCCGGACGTTAAAT-3’). A template for homologous recombination was designed to contain an HA-tag at the N-terminus of *Dicer* coding sequence and DNA sequence of exon 2 to exon 7 of Dicer (Fig. 1A). Detailed description of preparation of the template for homologous recombination template and its sequence are provided in the supplemental material.

Upon screening ∼600 ESC clones by PCR genotyping and Western blotting, we selected four ESC lines (Fig. S1), which were used to produce chimeric mice. *Dicer*^SOM^ ESCs were injected into eight-cell stage of C57Bl/6 host embryos. In total, 255 chimeric embryos were constructed, which were transferred into pseudopregnant foster mothers of ICR strain background. Of 27 weaned animals, fur color indicated chimerism in seven males and one female. Five males showing the strongest chimerism were mated to C57Bl/6 females to obtain germline transmission. From ∼ 140 genotyped progeny, we obtained one positive animal derived from the ESC clone 11 (Fig. S1), which was subsequently crossed on the C57Bl/6 background to establish the *Dicer*^SOM^ line.

### Oocyte collection

Fully-grown GV oocytes were obtained from superovulated or non-stimulated 12 – 16 weeks old C57Bl/6 mice as described previously [69]. Resumption of meiosis was prevented with 0.2 mM 3-isobutyl-1-methyl-xanthine (IBMX, Sigma). MII oocytes were obtained from animals superovulated with 7 IU of PMSG administered on day 1 and 7 IU of hCG administered 46 – 48 h post PMSG, mice were sacrificed 16 h post hCG. Oocytes were collected from oviducts, cumulus cells were removed by incubation with 0.1 uM hyaluronidase solution in M2 media for a 3 - 5 min, oocytes were washed from hyaluronidase in M2 media and collected in PBS-PVP.

### Cell culture and transfection

Mouse R1 ESCs were cultured in LIF media: KO-DMEM supplemented with 15% fetal calf serum, 1x L-Glutamine (Invitrogen), 1x non-essential amino acids (Invitrogen), 50 µM β-Mercaptoethanol (Gibco), 1000 U/mL LIF (ISOKine, ORF Genetics), penicillin (100 U/mL), and streptomycin (100 µg/mL). For transfection, cells were plated on a 24-well plate, grown to 50 % density and transfected using Lipofectamine 3000 (Thermo Fisher Scientific) according to the manufacturer’s protocol.

### Western blotting

Mouse tissues were homogenized mechanically in RIPA lysis buffer (20 mM HEPES (pH 7.8), 100 mM NaCl, 1 mM EDTA (pH 8.0), 0.5 % IGEPAL-25 %, 1 mM fresh DTT, 0.5 mM PMSF, 1 mM NaF, 0.2 mM Na_3_VO_4_, supplemented with 2x protease inhibitor cocktail set (Millipore)), centrifuged for 15 min, at 16 000 g, 4 °C and supernatant was used for protein electrophoresis. Protein concentration was measured by Bradford assay, 40 μg of protein was loaded per late. ESCs were grown in 6-well plates. Before collection, cells were washed with PBS and lysed in RIPA lysis buffer with inhibitors.

Proteins were separated on 5.5% polyacrylamide gel and transferred on PVDF membrane (Millipore) using semi-dry blotting for 50 min, 35 V. The membrane was blocked in 5% skim milk in TTBS, Dicer was detected using rat antiHA 3F10 monoclonal primary antibody (Sigma-Aldrich) diluted 1:2 500, membrane was incubated overnight at 4 °C in blocking solution. Washed in TTBS buffer, incubated with secondary antibody (HRP-antiRat diluted 1:50 000 in TTBS). Incubated for 1h at room temperature. Washed in TTBS, signal was detected using Supersignal west femto substrate (Thermo Scientific). For tubulin detection, samples were run on 10% PAA gel and incubated with anti-Tubulin (Sigma, #T6074) mouse primary antibody diluted 1:5 000 and anti-mouse-HRP secondary antibody 1:50 000.

### qPCR analyses

Ten oocytes per one reverse transcription (RT) reaction were collected in 1 ul PBS-PVP. 20 U of Ribolock RI (Thermofisher), 1 ug of yeast total RNA (carrier RNA) and RNase free water up to 5 ul was added and oocytes were incubated at 85 °C for 5 min for lysis. Crude lysate was used for RT with Maxima H Minus Reverse Transcriptase (Thermo Scientific). Ten oocytes from the same mouse were used for a control RT minus reaction. cDNA equivalent of a half of oocyte was used per a qPCR reaction. Maxima SYBR Green qPCR Master Mix (Thermo Scientific) was used for qPCR.

### Histology

Ovaries were fixed in Bouin solution for 16 - 20 h. Next, they were washed 3x 5 min with PBS, 30 min in 50% EtOH and transferred in 70% EtOH (all steps at 4 C). Whole ovaries were dehydrated, cleared and perfused with molten paraffin during histological processing. Ovaries were then embedded in paraffin blocks and sectioned to 8 µm sections and stained with conventional hematoxylin eosin stain. Testes were fixed in Davidson’s solution for 16 – 20 h and processed the same way as ovaries. 6 µm tissue sections were used for Dicer detection.

### Immunofluorescent staining

For analyzing microtubule defects, oocytes were fixed for 1h at room temperature in 4% PFA, permeabilized in 0.1% Triton X-100 for 10 min at room temperature (RT), washed in 2% BSA, 0.01% Tween-20 in PBS for 3×10 min, Blocked in 2% BSA, 0.01% Tween-20 in PBS for 1h, stained by primary antibody anti-β-Tubulin-Cy3 (TUB 2.1) (Abcam, ab11309) 1:200 for 1 h at room tempertaure, washed 3x 10 min in 0.01% Tween-20 in PBS, stained by 1 μg/ml DAPI for 5 min, mounted in Vectashield (Vector laboratories). Images of oocytes were acquired on DM6000 upright wide field microscope, images were processed by ImageJ.

For detection of HA-tagged Dicer, slides with 6 μm testis sections were deparaffinized and subjected to antigen retrieval in citrate buffer (pH=6). Sections were then permeabilized with 0.1% Triton X-100 for 15 min, blocked in 10% normal donkey serum, 0.1M glycine, 2% BSA for 1h at room tempertaure, incubated with primary antibody in blocking buffer (polyclonal rabbit anti-HA antibody (Cell signaling, 1:100) overnight at 4 °C, incubation with secondary antibody in blocking buffer for 1 h at room temperature in the dark (donkey anti rabbit antibody conjugated with Alexa 488) diluted 1:800. Nuclei were stained with 1 μg/ml DAPI for 5 min, slides were mounted in Vectashield media (Vector labs).

### RNA sequencing

Total RNA was extracted from 25 GV oocytes using Arcturus Picopure RNA isolation kit according to the manufacturer’s protocol. RNA was eluted in 11 μl of RNAse free water. Integrity of RNA was confirmed using 2100 Bioanalyzer (Agilent Technologies). NGS libraries were generated using V2 version of Ovation RNA-seq kit from Nugen. Libraries were prepared according to the manufacturer’s protocol. cDNA fragmentation was performed on Bioruptor sonication device (Diagenode) as follows: 45 s ON, 30 s OFF for 21 cycles on low intensity, 100 ng fragmented cDNA was used for library preparation. Libraries were amplified by 9 cycles of PCR and quantified by Qubit HS DNA kit, their quality was assessed using 2100 Bioanalyzer HS DNA chip. RNAseq libraries were pooled and sequenced by 50 nt single-end reading using the Illumina HiSeq2000 platform at the Genomics Core Facility at EMBL. RNA-seq data were deposited in the Gene Expression Omnibus database under accession ID GSE132121.

### Bioinformatic analyses

#### RNA-seq mapping and expression analysis

All RNA-seq data were mapped onto indexed genome using STAR 2.5.3a [70]: *STAR -- readFilesIn ${FILE}.fastq.gz --genomeDir $REF_GENOME_INDEX --runThreadN 8 -- genomeLoad LoadAndRemove –limitBAMsortRAM 20000000000 --readFilesCommand unpigz – c --outFileNamePrefix ${FILE}. --outSAMtype BAM SortedByCoordinate --outReadsUnmapped Fastx --outFilterMultimapNmax 99999 --outFilterMismatchNoverLmax 0.2 --sjdbScore 2*

Read files were mapped onto mouse genome version mm10/GRCm38. Mapped reads were counted using program featureCounts [71]: *featureCounts -a GENCODE.M15 -F ${FILE} - -minOverlap 10 --fracOverlap 0.00 -s 0 -M -O --fraction -J -T 8 ${FILE}.bam*

The GENCODE gene set (GENCODE M15) was used for the annotation. The count values were normalized by *rlog* function from program suite DESeq2 [72], which normalizes the data to the library size. The values of log2-fold-change were calculated among all the replicates of the corresponding conditions. The median was reported as the final value. Data were visualized as scatterplots.

#### Correlation matrices

Correlation matrices were calculated on the same count values normalized to per kilobase per million (RPKM) units. The retrotransposons were annotated by RepeatMasker, the mapped reads were counted for the GROUP category defined in *RepeatMasker-retrotransposon-groups.csv* (Supplementary file 1). The values were normalized to RPKM.

Small RNA-seq data were mapped onto indexed genome using STAR 2.5.3a: *STAR -- readFilesIn ${FILE}.fastq.gz --genomeDir $REF_GENOME_INDEX --runThreadN 8 -- genomeLoad LoadAndRemove --limitBAMsortRAM 20000000000 --readFilesCommand unpigz - c --outFileNamePrefix ${FILE}. --outSAMtype BAM SortedByCoordinate --outReadsUnmapped Fastx --outFilterMismatchNmax 1 --outFilterMismatchNoverLmax 1 -- outFilterMismatchNoverReadLmax 1 --outFilterMatchNmin 16 --outFilterMatchNminOverLread 0 --outFilterScoreMinOverLread 0 --outFilterMultimapNmax 9999 --alignIntronMax 1 -- alignSJDBoverhangMin 999999999999*

The histograms of the read lengths were calculated on the primary read alignments for the GROUP category defined in *RepeatMasker-retrotransposon-groups.csv* (Supplementary file 1). The potential siRNA targets (Supplementary file 2) were manually curated using the small RNA-seq datasets [32, 41, 43] selecting annotated protein-coding genes for which existed antisense endogenous 21-23nt small RNAs from traceable genomic origin, which would have over 50 RPM average abundance.

#### Retrotransposon expression analysis

For retrotransposon expression heatmap in germline (Fig. 5A), expression data from four different published datasets were mapped onto genome as described above GSE35005 [56]; GSE70116 [45], GSE45719 [57], and GSE41908 [58]. Next, reads overlapping one of the chosen retrotransposon subfamilies annotated by RepeatMasker [12] were counted with the following restrictions: (1) only perfectly mapping reads (i. e. nM tag in SAM files = 0) were counted, (2) only reads completely overlapping retrotransposon coordinates were counted, (3) coordinates of retrotransposons overlapping with genes annotation from Ensembl database (version 93) were removed from counting, and (4) each sequencing read mapped to more than one genomic sequence belonging to the same retrotransposon subfamily was counted only once. Count values were then normalized for library size using number of perfectly mapping reads in millions to obtain RPM values. For experiments with multiple replicates the RPM was calculated as the mean RPM value over all replicates.

In order to examine expression of 2691 full-length or nearly full-length L1 elements with ORF1 and ORF2 coding sequences of expected lengths (Fig. 5B; supplementary file 3) in fully-grown GV oocytes (125 PE from GSE116771 [62], 50 SE from this work), RNA-seq reads were mapped onto genome with all sequences except the 2691 L1 elements masked. Only reads perfectly mapping to those sequences were retained.

## ACKNOWLEDGEMENTS

We thank Vladimir Benes and EMBL sequencing facility for help with RNA-seq experiments, Kristian Vlahovicek (Zagreb University, Coratia) for providing hardware support for bioinformatics analysis, Donal O’Carroll (MRC Centre for Regenerative Medicine, University of Edinburgh, UK) for providing *Mili*^DAH^ mice. This work was funded from the European Research Council under the European Union’s Horizon 2020 research and innovation programme (grant agreement No 647403, D-FENS). Additional support was provided by the Ministry of Education, Youth, and Sports (MEYS) project NPU1 LO1419 (Biomodels for health). Financial support of E.T. and F.H. was in part provided by the Charles University through a PhD student fellowship; this work will be in part used to fulfill requirements for a PhD degree and hence can be considered “school work”. FH and ZJM were supported by the European Structural and Investment Funds grant for the Croatian National Centre of Research Excellence in Personalized Healthcare (contract #KK.01.1.1.01.0010), Croatian National Centre of Research Excellence for Data Science and Advanced Cooperative Systems (contract #KK.01.1.1.01.0009) and Croatian Science Foundation (grant IP-2014-09-6400). Production and histology analysis of the *Dicer*^SOM^ mice was supported by RVO 68378050 by Academy of Sciences of the Czech Republic and by LM2015040 (Czech Centre for Phenogenomics), CZ.1.05/2.1.00/19.0395 (Higher quality and capacity for transgenic models), CZ.1.05/1.1.00/02.0109 (BIOCEV - Biotechnology and Biomedicine Centre of the Academy of Sciences and Charles University). Microscopy was done at the Light Microscopy Core Facility, IMG CAS, Prague, Czech Republic supported by LM2015062 (Czech-Bioimaging). Additional computational resources were provided by CESNET LM2015042.

## SUPPORTING INFORMATION LEGENDS

**Figure S1** Analysis of ESC lines for *Dicer^SOM^* mouse model production by PCR and Western blotting. The yellow marked ESC lines were used for producing chimeric mice, the clone 11 (G2) gave rise to *Dicer^SOM^* animals used in the experiment.

**Figure S2** Comparison of transcriptomes of oocytes affected by different mutations in RNAi pathway. (A) Pairwise comparisons of relative gene expression changes using DESeq2-normalized expression values. In red are depicted genes potentially targeted by RNAi (i.e. with identified antisense siRNAs). (B) Correlation matrices calculated from RPKM values all annotated genes (upper panel) and predicted siRNA targets (lower panel).

**Supplementary Methods and TableS1 (Primer Table)**

**Supplementary files for bioinformatics analyses**

*file 1 RepeatMasker retrotransposon groups*

*file 2 siRNA target list*

*file 3 L1 table*

## REFERENCES

1. Bourque G, Burns KH, Gehring M, Gorbunova V, Seluanov A, Hammell M, et al. Ten things you should know about transposable elements. Genome Biol. 2018;19(1):199. doi: 10.1186/s13059-018-1577-z. PubMed PMID: 30454069; PubMed Central PMCID: PMCPMC6240941.

2. Craig NL, Chandler M, Gellert M, Lambowitz AM, Rice PA, Sandmeyer SB. Mobile DNA III. Craig NL, editor: AMS press; 2015.

3. International Human Genome Sequencing Consortium, Lander ES, Linton LM, Birren B, Nusbaum C, Zody MC, et al. Initial sequencing and analysis of the human genome. Nature. 2001;409(6822):860–921. doi: 10.1038/35057062. PubMed PMID: 11237011.

4. Mouse Genome Sequencing Consortium, Waterston RH, Lindblad-Toh K, Birney E, Rogers J, Abril JF, et al. Initial sequencing and comparative analysis of the mouse genome. Nature. 2002;420(6915):520–62. doi: 10.1038/nature01262. PubMed PMID: 12466850.

5. Kazazian HH, Jr. Mobile elements and disease. Curr Opin Genet Dev. 1998;8(3):343–50. PubMed PMID: 9690999.

6. Kuff EL, Lueders KK. The intracisternal A-particle gene family: structure and functional aspects. Adv Cancer Res. 1988;51:183–276. PubMed PMID: 3146900.

7. Dewannieux M, Dupressoir A, Harper F, Pierron G, Heidmann T. Identification of autonomous IAP LTR retrotransposons mobile in mammalian cells. Nat Genet. 2004;36(5):534–9. doi: 10.1038/ng1353. PubMed PMID: 15107856.

8. Ostertag EM, Kazazian HH, Jr. Biology of mammalian L1 retrotransposons. Annu Rev Genet. 2001;35:501–38. doi: 10.1146/annurev.genet.35.102401.091032. PubMed PMID: 11700292.

9. DeBerardinis RJ, Kazazian HH, Jr. Analysis of the promoter from an expanding mouse retrotransposon subfamily. Genomics. 1999;56(3):317–23. doi: 10.1006/geno.1998.5729. PubMed PMID: 10087199.

10. Speek M. Antisense promoter of human L1 retrotransposon drives transcription of adjacent cellular genes. Mol Cell Biol. 2001;21(6):1973–85. doi: 10.1128/MCB.21.6.1973-1985.2001. PubMed PMID: 11238933; PubMed Central PMCID: PMCPMC86790.

11. Wei W, Gilbert N, Ooi SL, Lawler JF, Ostertag EM, Kazazian HH, et al. Human L1 retrotransposition: cis preference versus trans complementation. Mol Cell Biol. 2001;21(4):1429–39. doi: 10.1128/MCB.21.4.1429-1439.2001. PubMed PMID: 11158327; PubMed Central PMCID: PMCPMC99594.

12. Smit AFA, Hubley R, Green P. RepeatMasker Open-4.0. <http://www.repeatmasker.org>.. 2013-2015.

13. Crichton JH, Dunican DS, Maclennan M, Meehan RR, Adams IR. Defending the genome from the enemy within: mechanisms of retrotransposon suppression in the mouse germline. Cell Mol Life Sci. 2014;71(9):1581–605. doi: 10.1007/s00018-013-1468-0. PubMed PMID: 24045705; PubMed Central PMCID: PMC3983883.

14. Ernst C, Odom DT, Kutter C. The emergence of piRNAs against transposon invasion to preserve mammalian genome integrity. Nat Commun. 2017;8(1):1411. doi: 10.1038/s41467-017-01049-7. PubMed PMID: 29127279; PubMed Central PMCID: PMC5681665.

15. Brennecke J, Aravin AA, Stark A, Dus M, Kellis M, Sachidanandam R, et al. Discrete small RNA-generating loci as master regulators of transposon activity in Drosophila. Cell. 2007;128(6):1089–103. doi: 10.1016/j.cell.2007.01.043. PubMed PMID: WOS:000245396200016.

16. Roovers EF, Rosenkranz D, Mahdipour M, Han CT, He N, Chuva de Sousa Lopes SM, et al. Piwi proteins and piRNAs in mammalian oocytes and early embryos. Cell Rep. 2015;10(12):2069–82. doi: 10.1016/j.celrep.2015.02.062. PubMed PMID: 25818294.

17. Kuramochi-Miyagawa S, Kimura T, Ijiri TW, Isobe T, Asada N, Fujita Y, et al. Mili, a mammalian member of piwi family gene, is essential for spermatogenesis. Development. 2004;131(4):839–49. doi: 10.1242/dev.00973. PubMed PMID: WOS:000189359500014.

18. Deng W, Lin HF. miwi, a murine homolog of piwi, encodes a cytoplasmic protein essential for spermatogenesis. Developmental Cell. 2002;2(6):819–30. doi: 10.1016/s1534-5807(02)00165-x. PubMed PMID: WOS:000176164000017.

19. Carmell MA, Girard A, van de Kant HJG, Bourc’his D, Bestor TH, de Rooij DG, et al. MIWI2 is essential for spermatogenesis and repression of transposons in the mouse male germline. Developmental Cell. 2007;12(4):503–14. doi: 10.1016/j.devcel.2007.03.001. PubMed PMID: WOS:000245816100007.

20. Lim AK, Lorthongpanich C, Chew TG, Tan CW, Shue YT, Balu S, et al. The nuage mediates retrotransposon silencing in mouse primordial ovarian follicles. Development. 2013;140(18):3819–25. Epub 2013/08/09. doi: 10.1242/dev.099184. PubMed PMID: 23924633; PubMed Central PMCID: PMCPMC4067262.

21. Kabayama Y, Toh H, Katanaya A, Sakurai T, Chuma S, Kuramochi-Miyagawa S, et al. Roles of MIWI, MILI and PLD6 in small RNA regulation in mouse growing oocytes. Nucleic Acids Research. 2017;45(9):5387–98. doi: 10.1093/nar/gkx027. PubMed PMID: WOS:000402064200040.

22. Aravin AA, Sachidanandam R, Bourc’his D, Schaefer C, Pezic D, Toth KF, et al. A piRNA pathway primed by individual transposons is linked to de novo DNA methylation in mice. Molecular Cell. 2008;31(6):785–99. doi: 10.1016/j.molcel.2008.09.003. PubMed PMID: WOS:000259721600004.

23. Kuramochi-Miyagawa S, Watanabe T, Gotoh K, Totoki Y, Toyoda A, Ikawa M, et al. DNA methylation of retrotransposon genes is regulated by Piwi family members MILI and MIWI2 in murine fetal testes. Genes Dev. 2008;22(7):908–17. Epub 2008/04/03. doi: 10.1101/gad.1640708. PubMed PMID: 18381894; PubMed Central PMCID: PMCPMC2279202.

24. De Fazio S, Bartonicek N, Di Giacomo M, Abreu-Goodger C, Sankar A, Funaya C, et al. The endonuclease activity of Mili fuels piRNA amplification that silences LINE1 elements. Nature. 2011;480(7376):259–63. doi: 10.1038/nature10547. PubMed PMID: 22020280.

25. Watanabe T, Cui X, Yuan Z, Qi H, Lin H. MIWI2 targets RNAs transcribed from piRNA-dependent regions to drive DNA methylation in mouse prospermatogonia. EMBO J. 2018;37(18). doi: 10.15252/embj.201695329. PubMed PMID: 30108053; PubMed Central PMCID: PMCPMC6138435.

26. Pezic D, Manakov SA, Sachidanandam R, Aravin AA. piRNA pathway targets active LINE1 elements to establish the repressive H3K9me3 mark in germ cells. Genes & Development. 2014;28(13):1410–28. doi: 10.1101/gad.240895.114. PubMed PMID: WOS:000338816000004.

27. Di Giacomo M, Comazzetto S, Saini H, De Fazio S, Carrieri C, Morgan M, et al. Multiple Epigenetic Mechanisms and the piRNA Pathway Enforce LINE1 Silencing during Adult Spermatogenesis. Molecular Cell. 2013;50(4):601–8. doi: 10.1016/j.molcel.2013.04.026. PubMed PMID: WOS:000319893600014.

28. Manakov S, Pezic D, Marinov G, Pastor W, Sachidanandam R, Aravin A. MIWI2 and MILI Have Differential Effects on piRNA Biogenesis and DNA Methylation. Cell Reports. 2015;12(8):1234–43. doi: 10.1016/j.celrep.2015.07.036. PubMed PMID: WOS:000360182200003.

29. Molaro A, Falciatori I, Hodges E, Aravin AA, Marran K, Rafii S, et al. Two waves of de novo methylation during mouse germ cell development. Genes & Development. 2014;28(14):1544–9. doi: 10.1101/gad.244350.114. PubMed PMID: WOS:000339166100003.

30. Zheng K, Xiol J, Reuter M, Eckardt S, Leu NA, McLaughlin KJ, et al. Mouse MOV10L1 associates with Piwi proteins and is an essential component of the Piwi-interacting RNA (piRNA) pathway. Proceedings of the National Academy of Sciences of the United States of America. 2010;107(26):11841–6. doi: 10.1073/pnas.1003953107. PubMed PMID: WOS:000279332300035.

31. Stein P, Rozhkov NV, Li F, Cardenas FL, Davydenko O, Vandivier LE, et al. Essential Role for endogenous siRNAs during meiosis in mouse oocytes. PLoS Genet. 2015;11(2):e1005013. doi: 10.1371/journal.pgen.1005013. PubMed PMID: 25695507; PubMed Central PMCID: PMCPMC4335007.

32. Tam OH, Aravin AA, Stein P, Girard A, Murchison EP, Cheloufi S, et al. Pseudogene-derived small interfering RNAs regulate gene expression in mouse oocytes. Nature. 2008;453(7194):534–8. Epub 2008/04/12. doi: 10.1038/nature06904. PubMed PMID: 18404147; PubMed Central PMCID: PMCPMC2981145.

33. Fire A, Xu S, Montgomery MK, Kostas SA, Driver SE, Mello CC. Potent and specific genetic interference by double-stranded RNA in Caenorhabditis elegans. Nature. 1998;391(6669):806–11. PubMed PMID: 9486653.

34. Wianny F, Zernicka-Goetz M. Specific interference with gene function by double-stranded RNA in early mouse development. Nat Cell Biol. 2000;2(2):70–5.

35. Svoboda P, Stain P, Hayashi H, Schultz RM. Selective reduction of dormant maternal mRNAs in mouse oocytes by RNA interference. Development. 2000;127(19):4147–56. PubMed PMID: WOS:000165122000007.

36. Svoboda P. Renaissance of mammalian endogenous RNAi. FEBS Lett. 2014;588(15):2550–6. doi: 10.1016/j.febslet.2014.05.030. PubMed PMID: 24873877.

37. Svoboda P, Stein P, Anger M, Bernstein E, Hannon GJ, Schultz RM. RNAi and expression of retrotransposons MuERV-L and IAP in preimplantation mouse embryos. Dev Biol. 2004;269(1):276–85. doi: 10.1016/j.ydbio.2004.01.028. PubMed PMID: 15081373.

38. Murchison EP, Stein P, Xuan Z, Pan H, Zhang MQ, Schultz RM, et al. Critical roles for Dicer in the female germline. Genes Dev. 2007;21(6):682–93. PubMed PMID: 17369401.

39. Tang F, Kaneda M, O’Carroll D, Hajkova P, Barton SC, Sun YA, et al. Maternal microRNAs are essential for mouse zygotic development. Genes Dev. 2007;21(6):644–8. PubMed PMID: 17369397.

40. Watanabe T, Totoki Y, Toyoda A, Kaneda M, Kuramochi-Miyagawa S, Obata Y, et al. Endogenous siRNAs from naturally formed dsRNAs regulate transcripts in mouse oocytes. Nature. 2008;453(7194):539–43. Epub 2008/04/12. doi: 10.1038/nature06908. PubMed PMID: 18404146.

41. Garcia-Lopez J, Hourcade Jde D, Alonso L, Cardenas DB, del Mazo J. Global characterization and target identification of piRNAs and endo-siRNAs in mouse gametes and zygotes. Biochim Biophys Acta. 2014;1839(6):463–75. doi: 10.1016/j.bbagrm.2014.04.006. PubMed PMID: 24769224.

42. Larriba E, Del Mazo J. An integrative piRNA analysis of mouse gametes and zygotes reveals new potential origins and gene regulatory roles. Sci Rep. 2018;8(1):12832. Epub 2018/08/29. doi: 10.1038/s41598-018-31032-1. PubMed PMID: 30150632; PubMed Central PMCID: PMCPMC6110870.

43. Yang Q, Lin J, Liu M, Li R, Tian B, Zhang X, et al. Highly sensitive sequencing reveals dynamic modifications and activities of small RNAs in mouse oocytes and early embryos. Sci Adv. 2016;2(6):e1501482. doi: 10.1126/sciadv.1501482. PubMed PMID: 27500274; PubMed Central PMCID: PMCPMC4974095.

44. Flemr M, Malik R, Franke V, Nejepinska J, Sedlacek R, Vlahovicek K, et al. A retrotransposon-driven dicer isoform directs endogenous small interfering RNA production in mouse oocytes. Cell. 2013;155(4):807–16. Epub 2013/11/12. doi: 10.1016/j.cell.2013.10.001. PubMed PMID: 24209619.

45. Veselovska L, Smallwood SA, Saadeh H, Stewart KR, Krueger F, Maupetit-Mehouas S, et al. Deep sequencing and de novo assembly of the mouse oocyte transcriptome define the contribution of transcription to the DNA methylation landscape. Genome Biol. 2015;16:209. doi: 10.1186/s13059-015-0769-z. PubMed PMID: 26408185; PubMed Central PMCID: PMC4582738.

46. Franke V, Ganesh S, Karlic R, Malik R, Pasulka J, Horvat F, et al. Long terminal repeats power evolution of genes and gene expression programs in mammalian oocytes and zygotes. Genome Research. 2017;27(8):1384–94. doi: 10.1101/gr.216150.116. PubMed PMID: WOS:000406354300009.

47. Much C, Auchynnikava T, Pavlinic D, Buness A, Rappsilber J, Benes V, et al. Endogenous Mouse Dicer Is an Exclusively Cytoplasmic Protein. Plos Genetics. 2016;12(6). doi: 10.1371/journal.pgen.1006095. PubMed PMID: WOS:000379347100015.

48. Houwing S, Kamminga LM, Berezikov E, Cronembold D, Girard A, van den Elst H, et al. A role for Piwi and piRNAs in germ cell maintenance and transposon silencing in zebrafish. Cell. 2007;129(1):69–82. doi: 10.1016/j.cell.2007.03.026. PubMed PMID: WOS:000245661100012.

49. Nejepinska J, Malik R, Filkowski J, Flemr M, Filipowicz W, Svoboda P. dsRNA expression in the mouse elicits RNAi in oocytes and low adenosine deamination in somatic cells. Nucleic Acids Research. 2012;40(1):399–413. doi: 10.1093/nar/gkr702. PubMed PMID: WOS:000298733500043.

50. Demeter T, Vaskovicova M, Malik R, Horvat F, Pasulka J, Svobodova E, et al. Main constraints for RNAi induced by expressed long dsRNA in mouse cells. Life Sci Alliance. 2019;2(1). doi: 10.26508/lsa.201800289. PubMed PMID: 30808654; PubMed Central PMCID: PMCPMC6391682.

51. Yang ZL, Chen KM, Pandey RR, Homolka D, Reuter M, Janeiro BKR, et al. PIWI Slicing and EXD1 Drive Biogenesis of Nuclear piRNAs from Cytosolic Targets of the Mouse piRNA Pathway. Molecular Cell. 2016;61(1):138–52. doi: 10.1016/j.molcel.2015.11.009. PubMed PMID: WOS:000372324500012.

52. Zheng K, Wang PJ. Blockade of pachytene piRNA biogenesis reveals a novel requirement for maintaining post-meiotic germline genome integrity. PLoS Genet. 2012;8(11):e1003038. doi: 10.1371/journal.pgen.1003038. PubMed PMID: 23166510; PubMed Central PMCID: PMCPMC3499362.

53. Edson MA, Nagaraja AK, Matzuk MM. The mammalian ovary from genesis to revelation. Endocr Rev. 2009;30(6):624–712. doi: 10.1210/er.2009-0012. PubMed PMID: 19776209; PubMed Central PMCID: PMCPMC2761115.

54. Aravin AA, Hannon GJ, Brennecke J. The Piwi-piRNA pathway provides an adaptive defense in the transposon arms race. Science. 2007;318(5851):761–4. doi: 10.1126/science.1146484. PubMed PMID: WOS:000250583900031.

55. Horman SR, Svoboda P, Luning Prak ET. The potential regulation of L1 mobility by RNA interference. J Biomed Biotechnol. 2006;2006(1):32713. doi: 10.1155/JBB/2006/32713. PubMed PMID: 16877813; PubMed Central PMCID: PMCPMC1559915.

56. Gan H, Cai T, Lin X, Wu Y, Wang X, Yang F, et al. Integrative proteomic and transcriptomic analyses reveal multiple post-transcriptional regulatory mechanisms of mouse spermatogenesis. Mol Cell Proteomics. 2013;12(5):1144–57. doi: 10.1074/mcp.M112.020123. PubMed PMID: 23325766; PubMed Central PMCID: PMCPMC3650327.

57. Deng Q, Ramskold D, Reinius B, Sandberg R. Single-cell RNA-seq reveals dynamic, random monoallelic gene expression in mammalian cells. Science. 2014;343(6167):193–6. doi: 10.1126/science.1245316. PubMed PMID: 24408435.

58. Yamaguchi S, Hong K, Liu R, Inoue A, Shen L, Zhang K, et al. Dynamics of 5-methylcytosine and 5-hydroxymethylcytosine during germ cell reprogramming. Cell Res. 2013;23(3):329–39. doi: 10.1038/cr.2013.22. PubMed PMID: 23399596; PubMed Central PMCID: PMCPMC3587712.

59. DeBerardinis RJ, Goodier JL, Ostertag EM, Kazazian HH, Jr. Rapid amplification of a retrotransposon subfamily is evolving the mouse genome. Nat Genet. 1998;20(3):288–90. doi: 10.1038/3104. PubMed PMID: 9806550.

60. Hardies SC, Wang L, Zhou L, Zhao Y, Casavant NC, Huang S. LINE-1 (L1) lineages in the mouse. Mol Biol Evol. 2000;17(4):616–28. doi: 10.1093/oxfordjournals.molbev.a026340. PubMed PMID: 10742052.

61. Goodier JL, Ostertag EM, Du K, Kazazian HH, Jr. A novel active L1 retrotransposon subfamily in the mouse. Genome Res. 2001;11(10):1677–85. doi: 10.1101/gr.198301. PubMed PMID: 11591644; PubMed Central PMCID: PMCPMC311137.

62. Horvat F, Fulka H, Jankele R, Malik R, Jun M, Solcova K, et al. Role of Cnot6l in maternal mRNA turnover. Life Sci Alliance. 2018;1(4):e201800084. doi: 10.26508/lsa.201800084. PubMed PMID: 30456367; PubMed Central PMCID: PMCPMC6238536.

63. Newkirk SJ, Lee S, Grandi FC, Gaysinskaya V, Rosser JM, Berg NV, et al. Intact piRNA pathway prevents L1 mobilization in male meiosis. Proceedings of the National Academy of Sciences of the United States of America. 2017;114(28):E5635–E44. doi: 10.1073/pnas.1701069114. PubMed PMID: WOS:000405177100020.

64. Malki S, van der Heijden GW, O’Donnell KA, Martin SL, Bortvin A. A Role for Retrotransposon LINE-1 in Fetal Oocyte Attrition in Mice. Developmental Cell. 2014;29(5):521–33. doi: 10.1016/j.devcel.2014.04.027. PubMed PMID: WOS:000337644700006.

65. Ecco G, Imbeault M, Trono D. KRAB zinc finger proteins. Development. 2017;144(15):2719–29. doi: 10.1242/dev.132605. PubMed PMID: 28765213.

66. Castro-Diaz N, Ecco G, Coluccio A, Kapopoulou A, Yazdanpanah B, Friedli M, et al. Evolutionally dynamic L1 regulation in embryonic stem cells. Genes Dev. 2014;28(13):1397–409. doi: 10.1101/gad.241661.114. PubMed PMID: 24939876; PubMed Central PMCID: PMCPMC4083085.

67. Nagy A, Rossant J, Nagy R, Abramow-Newerly W, Roder JC. Derivation of completely cell culture-derived mice from early-passage embryonic stem cells. Proc Natl Acad Sci U S A. 1993;90(18):8424–8. doi: 10.1073/pnas.90.18.8424. PubMed PMID: 8378314; PubMed Central PMCID: PMCPMC47369.

68. Ran FA, Hsu PD, Wright J, Agarwala V, Scott DA, Zhang F. Genome engineering using the CRISPR-Cas9 system. Nat Protoc. 2013;8(11):2281–308. doi: 10.1038/nprot.2013.143. PubMed PMID: 24157548; PubMed Central PMCID: PMCPMC3969860.

69. Nagy A. Manipulating the mouse embryo: a laboratory manual. 3rd ed. Cold Spring Harbor, N.Y.: Cold Spring Harbor Laboratory Press; 2003. x, 764 p. p.

70. Dobin A, Davis CA, Schlesinger F, Drenkow J, Zaleski C, Jha S, et al. STAR: ultrafast universal RNA-seq aligner. Bioinformatics. 2013;29(1):15–21. doi: 10.1093/bioinformatics/bts635. PubMed PMID: 23104886; PubMed Central PMCID: PMCPMC3530905.

71. Liao Y, Smyth GK, Shi W. featureCounts: an efficient general purpose program for assigning sequence reads to genomic features. Bioinformatics. 2014;30(7):923–30. doi: 10.1093/bioinformatics/btt656. PubMed PMID: 24227677.

72. Love MI, Huber W, Anders S. Moderated estimation of fold change and dispersion for RNA-seq data with DESeq2. Genome Biol. 2014;15(12):550. doi: 10.1186/s13059-014-0550-8. PubMed PMID: 25516281; PubMed Central PMCID: PMCPMC4302049.

73. Abe K, Yamamoto R, Franke V, Cao M, Suzuki Y, Suzuki MG, et al. The first murine zygotic transcription is promiscuous and uncoupled from splicing and 3’ processing. EMBO J. 2015;34(11):1523–37. doi: 10.15252/embj.201490648. PubMed PMID: 25896510; PubMed Central PMCID: PMCPMC4474528.

